# Endosomal proteins NPC1 and NPC2 at African swine fever virus entry/fusion

**DOI:** 10.1101/2021.07.07.451424

**Authors:** Miguel Ángel Cuesta-Geijo, Jesús Urquiza, Ana del Puerto, Isabel Garcia-Dorival, Fátima Lasala, Lucía Barrado-Gil, Inmaculada Galindo, Rafael Delgado, Covadonga Alonso

## Abstract

African swine fever virus (ASFV) infectious cycle starts with the viral adsorption and entry into the host cell. The virus is internalized via clathrin/dynamin mediated endocytosis and macropinocytosis. As several other viruses, ASF virion is then internalized and incorporated into the endocytic pathway. Endosomal maturation entails luminal acidification and the lowering of pH acting on the multi-layered virion structure dissolves the outer capsid. Upon decapsidation, the inner viral membrane is exposed to interact with the limiting membrane of the late endosome for fusion. Egress from endosome is related to cholesterol efflux, but it remains an intriguing process albeit essential for infection, specifically for the viral nucleic acid exit to the cytoplasm for replication. ASFV proteins E248R and E199L, with structural homology to the VACV proteins of the fusion complex, seem to have similar functions in ASFV. A direct interaction between these ASFV proteins with the cholesterol transporter protein NPC1 (Niemann-Pick C type 1) was observed, which was also shared by the E248R homologous protein L1R of VACV. Binding occurs between the transmembrane domain of E248R with the loop C of NPC1 at the same domain than EBOV binding site. These interactions suggest that these ASFV proteins are crucial for membrane fusion. CRISPR NPC1 KO Vero cells lacking NPC1 protein that were resistant to EBOV, reduced ASFV infection levels significantly. Reductions on ASFV infectivity and replication in NPC1 KO cells were accompanied by lesser viral factories of smaller size and lacking the typical cohesive morphology between endosomes and viral proteins. We observed a compensatory effect in NPC1 KO cells, elevating NPC2 levels while silencing NPC2 in Vero cells with shRNA, also reduced ASFV infection. Our findings pave the way to understand the role of these proteins at the membrane viral fusion step for several viruses.

**Author Summary:** African swine fever virus (ASFV) causes a deadly disease of pigs and wild boars that was endemic in Africa but have extended over the last years to Europe, Asia and Oceania with high socioeconomic impact. ASFV enters the cell by endocytosis and has adapted to the endosomal conditions to acquire infectivity. Viral infectivity is dependent on cholesterol traffic at the endosomes, especially at the fusion step. Fusion of the internal viral membrane with the endosomal membrane is required for the exit of the viral DNA to the cytoplasm to start replication. ASF virion internal membrane proteins E248R and E199L were found to bind the Niemann Pick C1 (NPC1) receptor at the endosome. These proteins are highly conserved among ASFV isolates and resemble proteins of the VACV entry/fusion complex. The function of NPC1 is to regulate the efflux of dietary cholesterol efflux from the endosome to the endoplasmic reticulum, which appears to be necessary for viral fusion. NPC1 knockout cells by CRISPR reduced infection affecting infectivity and early replication. Also, removing the associated endosomal protein NPC2, further declined infectivity. These results show the relevance of NPC1 receptor in the viral infection actually shared by unrelated important viral families.

## INTRODUCTION

African Swine Fever Virus (ASFV) is the only known member of the *Asfarviridae* family, and the only known DNA arbovirus. It is a large, enveloped virus with an average diameter of 200 nm and a multi-layered structure and icosahedral morphology that has been recently unveiled in detail (1–3).

ASFV is the causative agent of African Swine Fever (ASF), a high mortality haemorrhagic disease affecting swine which is endemic in sub-Saharian Africa. However, ASF epidemics that started in the Caucasus and Russian Federation in 2007 (4), have expanded rapidly over different countries in Europe, Asia and Oceania; causing a devastating burden in the global pig industry (5). Currently, ASF cases have been reported in Germany since September 2020 (6); furthermore, India has also recently declared an ASF outbreak (7, 8).

ASFV infectious cycle starts with the viral adsorption and entry into the host cell. After attachment to an unknown receptor, the virus is mainly internalized via clathrin/dynamin mediated endocytosis and micropinocytosis (9, 10). Then, the virion is internalized and incorporated into the endocytic pathway.

Under the molecular cues of endosomes, the multi-layered virion undergoes uncoating starting from decapsidation (11). This step will be followed by a less known fusion process at late endosomes (LE), delivering the naked viral core out to the cytoplasm.

According to (11), endosomal maturation entails dropping of luminal pH and this acidic environment disrupts the structure of the virion and produces decapsidation at 30-45 minutes post infection (mpi). The consequence is the exposure of the inner viral membrane allowing its fusion with the limiting membrane of the LE. This fusion is strongly dependent on cholesterol efflux at the LE. In fact, blocking cholesterol transport at this level causes retention of virions inside endosomes, and inhibits infection progression (12).

Cholesterol transport from LE requires the acidic pH (13) and it is mainly regulated by the coordinated action of LE proteins Niemann Pick type C 1 (NPC1), an endolysosomal membrane protein; and NPC2, a single domain protein located in the lumen of the vesicle (14–16).

Similar to other viruses, such as Ebola virus (EBOV) (17, 18), the ASFV replication cycle is strongly impacted by compounds such as U18666A or imipramine (19, 20). These compounds act either by binding to NPC1 sterol sensing domain (SSD) or by blocking EBOV Glycoprotein (GP) GP-NPC1 interaction, respectively. This indicates that NPC1 and intact cholesterol transport are pivotal for a successful infection (21–32).

Since the underlying mechanisms of viral fusion and egress to the cytoplasm still remain obscure, we hypothesized that ASFV could penetrate to the cytoplasm from LE in a NPC1-dependent manner via proteins belonging to its fusion machinery.

ASFV proteins potentially involved in fusion would be probably internal membrane proteins E248R and E199L. These are structural proteins, late synthesized, which have been involved in ASF fusion and viral core penetration (33–35), lacking a demonstrated interaction between them (34). As both proteins weakly resemble to different subunits of the fusion complex of poxvirus, this makes them appropriate candidates to analyse their function at the viral core penetration to the cytoplasm.

ASFV protein E248R is a type II late structural protein required for viral membrane fusion with limiting membrane of LE (33, 35). It belongs to a class of myristoylated proteins related to vaccinia virus (VACV) such as VACV L1R protein, which forms part of the VACV fusion complex. L1R contains several disulfide bonds and it is widely conserved in several families of Poxvirus (36, 37).

ASFV E199L is a transmembrane protein type I harboring disulfide bonds and together with E248R, have been postulated to be necessary for membrane fusion and core penetration to the cytoplasm (34). E199L resemble to three poxviral fusion machinery subunits, named A16, G9 and A26. In absence of E199L, ASFV infection drastically decreased and viral particles are retained into LE and lysosomes.

In this report, we have shown that E248R and E199L, two candidate proteins potentially involved in ASFV membrane process and located in the inner membrane of the viral particle, interact with the LE integral membrane protein NPC1. This novel finding supports an interaction of the internal membrane proteins E248R and E199L with NPC1 C domain, thus bringing to the forefront this cholesterol transporter as a crucial target to counteract the early steps of the infection.

## RESULTS

### Cholesterol accumulation is critical for ASFV infection

A first hint of the influence of cholesterol flux on ASFV infectivity was obtained by using the U18666A chemical, a compound that blocks the NPC1 cholesterol transporter. This compound reduces the cholesterol efflux from the endosome, which remains retained inside these vesicles and its accumulation resulted in the formation of large, dilated endosomes. We pre-treated Vero cells with increasing concentrations of the U18666A drug and labelled cholesterol with Filipin antibody. By these means, we correlated cholesterol retention with reductions in ASFV infectivity at the same drug concentrations. Fig 1A shows a characteristic accumulation of cholesterol in large vesicle-like clusters around the perinuclear area starting at a drug concentration of 0.5 µM. Likewise, Vero cells were pre-treated with similar concentrations of U18666A drug and infected with ASFV BPP30 GFP at a moi of 1 pfu/ml for 16 hpi. After this time, GFP expression was tested in a plate reader. Importantly, infection started to be impaired at the same concentrations that produced a cholesterol accumulation phenotype, suggesting that there could be a correlation between both facts (Fig 1B).

**Fig 1.**
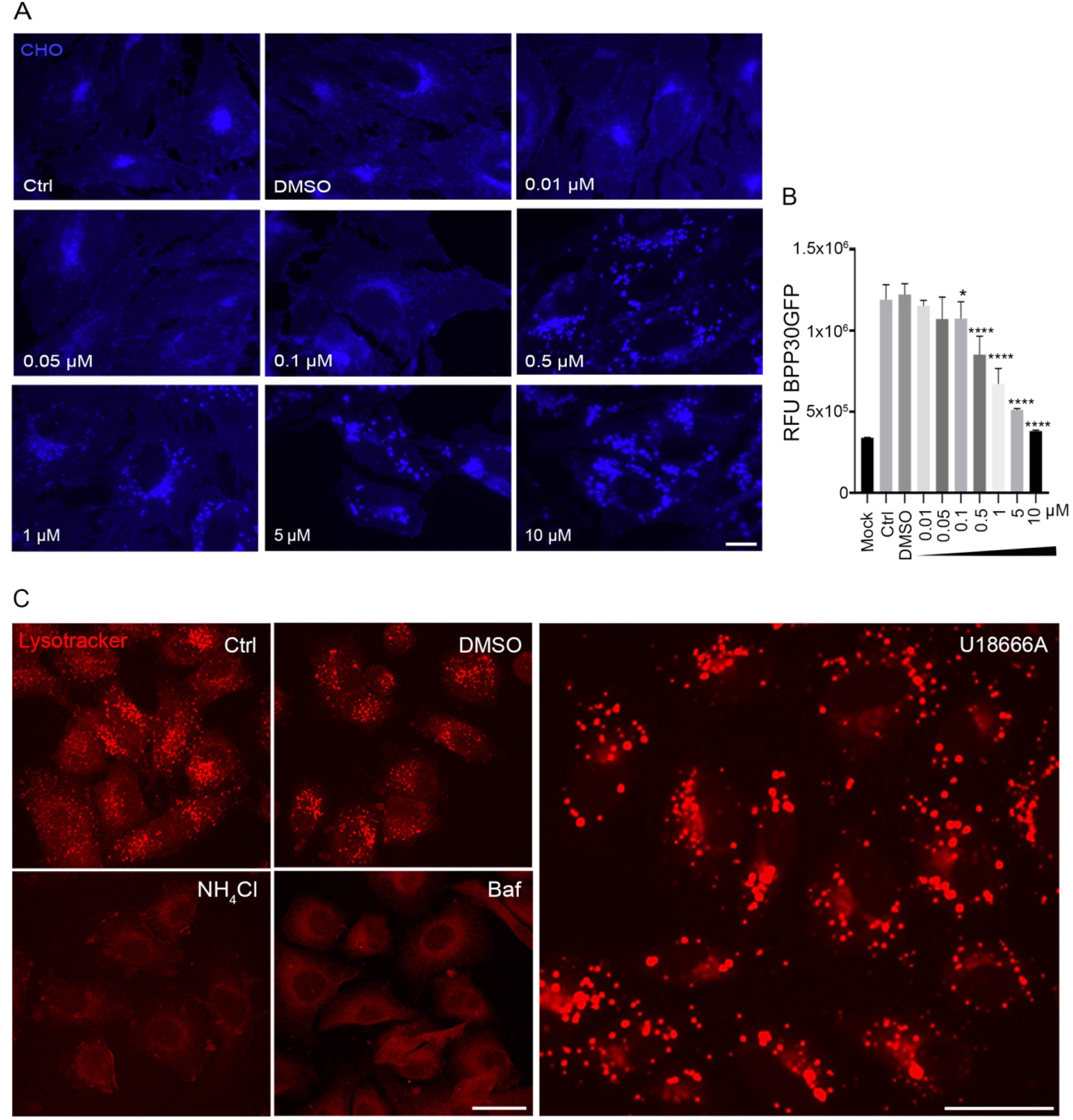
Cholesterol efflux blockade inhibition of ASFV infection. (A) Increasing concentrations of U18666A chemical compound altered the distribution of unesterified cholesterol (filipin; blue). Scale bar 10 µM. (B) Vero cells pre-treated at increasing concentrations of U drug were infected with recombinant ASFV PP30GFP. After 16 hpi, GFP fluorescence intensity was measured by plaque assay. (C) Visualization of acidic vesicles with lysotracker (red) in cells treated with U18666A drug or controls, but not after lysosomotropic drugs NH4Cl or Bafilomycin treatment. Scale bar 25µM.

Since ASFV is strongly dependent of acidic endosomal pH (11) we excluded the possibility of a lysosomotropic effect of the U drug by using the acid pH probe Lysotracker. As positive controls, Vero cells were pre-treated with lysosomotropic drugs Bafilomycin and NH_4_Cl. Acidic vesicles were clearly stained in controls and U18666A drug treated cells, while no signal from acidic vesicles was detected with Bafilomycin and NH_4_Cl. This confirmed that the U18666A compound acted specifically as an inhibitor of cholesterol export in this context and it has no action on acidification (Fig 1C).

The U18666A compound affects the cholesterol transporter function of NPC1 directly binding to the SSD of NPC1 resulting in the blockade of cholesterol efflux out of endolysosomes, which also inhibits EBOV infection (19). This suggested that the transporter role of NPC1 could be necessary for ASFV egress from the LE to the cytosol as it was described for EBOV. This would ultimately affect viral replication either if this occurs as a result of the lack of cholesterol efflux or because of a mechanistic impairment. In EBOV infection, the viral glycoprotein interacts directly with NPC1 to allow viral fusion and endosomal escape. Then, we set out to investigate a possible similar mechanism mediated by analyzing the direct binding of an ASFV protein with NPC1.

### E248R and E199L sequence analysis

As we mentioned above, E248R and E199L have been involved in membrane fusion to finally conduct the naked viral core of the endocytosed virions from the LE to the cytoplasm. These are late synthesized structural proteins highly conserved proteins among ASFV isolates (Fig 2A). Furthermore, both proteins resemble to different subunits of the entry/fusion multiprotein complex of poxvirus (S1 Fig).

**Fig 2.**
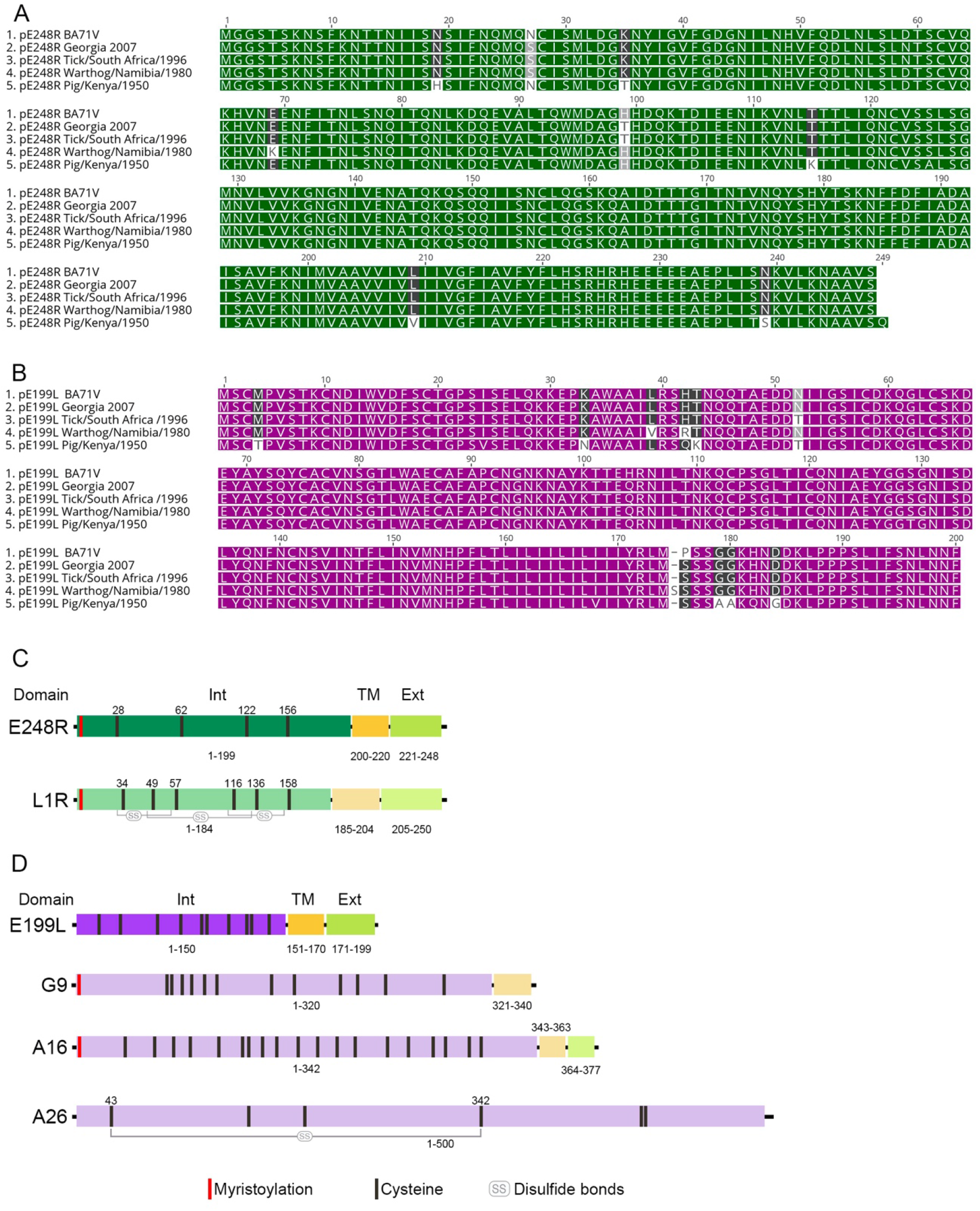
Comparison of ASFV E248R and E199L in isolates and with VACV fusion proteins main domains. (A-B) Identical amino acids are marked green for E248R or purple for E199L. Amino acids conserved 80% are marked in dark grey, those conserved between 60-80% in light grey. Finally, those preserved less than 60% in white. (C-D) Structural comparison with VACV fusion proteins. Myristoylation signal is indicated in red, the three main domains: internal (Int), transmembrane (TM) and external domains (Ext) represented in blocks. Amino acid numbers are indicated. Black lines represent cysteines indicating the positions above and the disulfide bonds in grey.

E248R is a type II myristoylated transmembrane protein. It is composed of an N-terminal portion of 199 amino acids, which contains four cysteine residues oriented to the cytoplasm, a helical hydrophobic transmembrane domain of 21 amino acids, and a 28 amino acid length extracellular region (35). E248R is the final substrate of the ASFV-encoded redox system (38) and, as a consequence, it could contain intramolecular disulfide bonds. We found that E248R yielded 16.2% identity and 30.7% similarity with VACV L1R fusion protein.

The distribution of cysteine residues in E248R (Fig 2C), compared to related protein of VACV protein L1R would suggest that the two L1R disulfide bonds between amino acids 28-62 and 122-156 could be also present in E248R (Fig. 2C and S1 Fig A).

E199L is a type I transmembrane protein with a N-terminal large cysteine enriched portion oriented internally to the viral particle, and small transmembrane and external (C-terminal) domains. E199L shares some degree of homology to three cysteine enriched proteins belonging to the fusion machinery of VACV named A16, A26 and G9, which also form disufide (39) bonds (Fig 2D and S1 Fig B).

In order to determine a functional mechanism that would relate both viral proteins with ASFV endosomal egress, we designed a protein-protein interaction experiment with NPC1, thus connecting cholesterol efflux in late endosomal vesicles with these proteins.

### Interaction of E248R and E199L protein with NPC1

Cholesterol efflux mediated by NPC1 is crucial in ASFV infection and NPC1 is an intracellular receptor for EBOV. Then, we tested the possibility of NPC1 being involved in the fusion of viral and LE membranes, along with the viral proteins E248R and E199L. These proteins located at the exposed inner membrane of the virion after decapsidation are candidates for membrane fusion and viral core penetration to the cytoplasm.

Schematics in Fig 3A depicts E248R domains and the deletion mutant constructions designed for the experiment. E248R ΔExt, lacking the external protein domain, E248R ΔTM obtained by deletion of the transmembrane domain, and E248R ΔExt+TM, obtained by deletion of both transmembrane and external domains. These deletions were constructed to clarify the relevance of each of the ASFV protein E248R domains, exposing their role for binding the cellular receptor.

**Fig 3.**
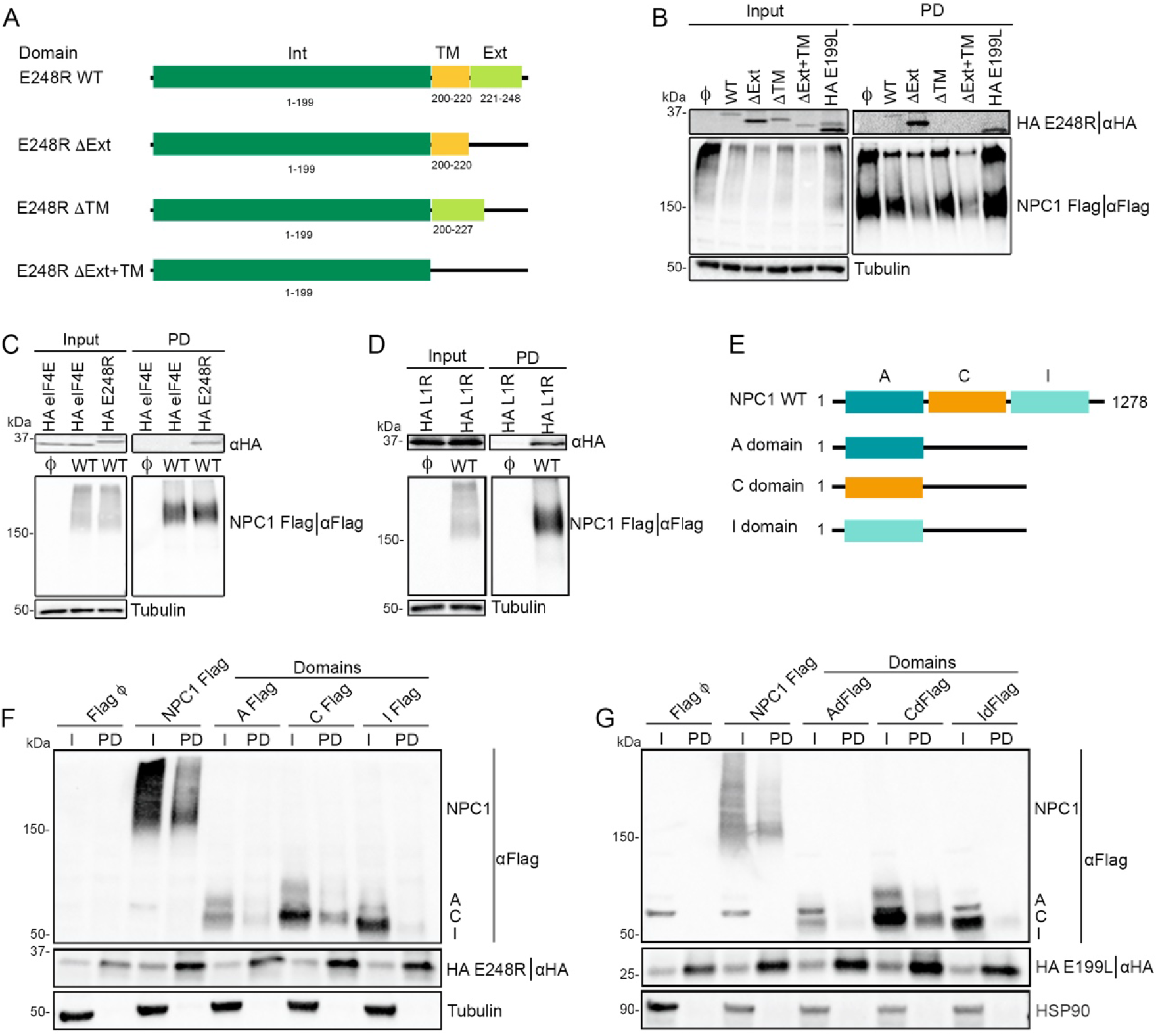
E248R and E199L interact with NPC1. (A) Schematic representation of E248R WT protein and deletion mutant domains. (B) Detection of HA E248R WT, HA E248R mutants and HA E199L WT co-expressed individually with NPC1 Flag in 293T cells. HA and Flag fused proteins analyzed in the immunoprecipitation assay (“PD” refers to pulled down sample) by western blot. (C) Flag pull-down experiments of 293T cells transiently co-expressing NPC1 Flag together with HA E248R or HA eIF4e as a negative control of the interaction analyzed by western blot. (D) Flag pull-down experiments of 293T cells transiently co-expressing NPC1 Flag together with HA L1R. (E) Schematic representation of NPC1 WT and constructions of individual domains. (F) Detection of co-expressed HA E248R with NPC1 Flag as a control for positive interaction, A Flag, C Flag or I Flag construction (“PD” is the pull down sample). Interactions were detected by reverse pulldown with Sepharose beads HA tagged were detected by western blot. (G) Detection of co-expressed HA E199L (as a positive control of interaction) with NPC1 Flag, A Flag, C Flag or I Flag construction (“PD” is the pull down sample). Interactions were detected by reverse pulldown with Sepharose beads HA tagged were detected by western blot. All immunoprecipitation assays experiments were repeated three times to ensure reproducibility.

First, to investigate the interaction of full-length E248R and E199L with NPC1-Flag, both viral proteins were expressed as HA-fusions in HEK 293T cells, and independently co-expressed with NPC1-Flag. HEK 293T cells were selected due to their high efficiency of transfection, being the cell line of choice for protein-protein interaction studies of several viruses.

Protein expression of HA-E248R, mutants, HA-E199L and NPC1-Flag was confirmed using western blot analysis (Fig 3B). NPC1-Flag or empty Flag were then co-immunoprecipitated using Flag beads. After immunoprecipitation, both input (cell lysate) and bound samples were analyzed by western blot.

Previous reports have shown the interaction between the primed form of GP glycoprotein of EBOV and NPC1 cholesterol transporter to promote fusion with the LE membrane (40). Since ASFV requires an intact cholesterol efflux from LE (12) and traffics to the cytoplasm from LE compartment (11, 33) similarly to EBOV, we analyse the putative implication of NPC1 in ASFV infection, through an interaction with the suspected ASFV proteins involved in fusion with the LE, E248R and E199L (33, 34).

As shown in Fig 3B, HA-E248R WT, HA-E248R mutants with TM domain, and HA-E199L were detected in the immunoprecipitated pellet with Flag beads with an HA antibody. NPC1 was also detected by using an antibody against Flag. Conversely, an independent cellular protein fused to HA (HA-eIF4E) used as a negative control was not detected (Fig 3C), corroborating the interaction specificity of E248R and E199L with NPC1. Altogether, these results demonstrate that E248R interacts through its TM domain with NPC1, in fact E248R mutants lacking TM domain shown no interaction with NPC1.

Simultaneously, we could also demonstrate VACV L1R direct binding to NPC1-Flag (Fig 3D).

### E248R and E199L interact with NPC1 through C domain

In order to identify the specific NPC1 domain involved in the interaction with E248R, we designed three constructions expressing NPC1 individual domains A, C, I fused to Flag (Fig 3E) (41). These domains were co-expressed individually with HA E248R for 24 hours in 293T cells. Co-immunoprecipitation with Flag beads was undetermined, depicting an interaction with the three different NPC1 domains A, C and I, as analyzed by western blot (data not shown).

To further characterize the specific interaction between E248R with NPC1 and the individual domains, the reciprocal co-immunoprecipitations (Co-IP or reverse pull downs) against HA E248R were performed using protein G-beads and specific monoclonal antibodies against HA (Fig 3F). The expression of HA E248R was confirmed using a HA antibody as expected but only NPC1 WT and C domain were retrieved in the bound samples obtained from the Co-IP and analyzed by WB.

A similar experiment was performed and resolved co-expressing HA E199L with NPC1 WT and A, C and I domains fussed to Flag. We found a clear interaction with NPC1 WT and C domain (Fig 3G).

Thus, this result confirms the interaction between E248R via TM domain and NPC1, through C domain of NPC1, similar to the EBOV GP interaction with NPC1 at the C domain. Also, E199L was found to interact with NPC1 via C domain.

### ASFV infection in NPC1 Knockout cells

After confirming the interaction of both proteins with NPC1, CRISPR Cas9 technology was used to generate a knockout (KO) Vero cell line for NPC1 expression. The absence of NPC1 was confirmed from the pool of knocked cells by western blot (WB), indirect immunofluorescence (IFI) and (S6 Fig A-B). Further assays were performed to test the cellular phenotype that should be compatible with the absence of NPC1 in the cells. The absence of NPC1 would result in a cholesterol accumulation phenotype in dilated acidic vesicles, as we observed in S6 Fig C. A similar cholesterol accumulation pattern than cells pre-treated with U18666A compound was also observed in NPC1 KO pool of cells (S6 Fig D). Moreover, since NPC1 acts as an intracellular receptor for filovirus, these KO cells should be resistant to filovirus infection, as we verify by inhibition of VSV-pseudotyped with GP EBOV Mayinga strain infection, confirming a functional abrogation of NPC1 (S6 Fig E).

Susceptibility to ASFV infection remained similar between parental Vero WT cells and a selected LCV_2_ clone (named as Empty), both were used as controls for infection of NPC1 KO cells. After re-confirming the KO of NPC1 protein expression by WB in several clones from the pool of KO cells, clone 14 (c14) (named as NPC1 KO) was selected to perform further experiments. Also, we stated the absence of NPC1 mRNA and the lack of protein expression by IFI (Fig 4A, 4B and 4C).

**Fig 4.**
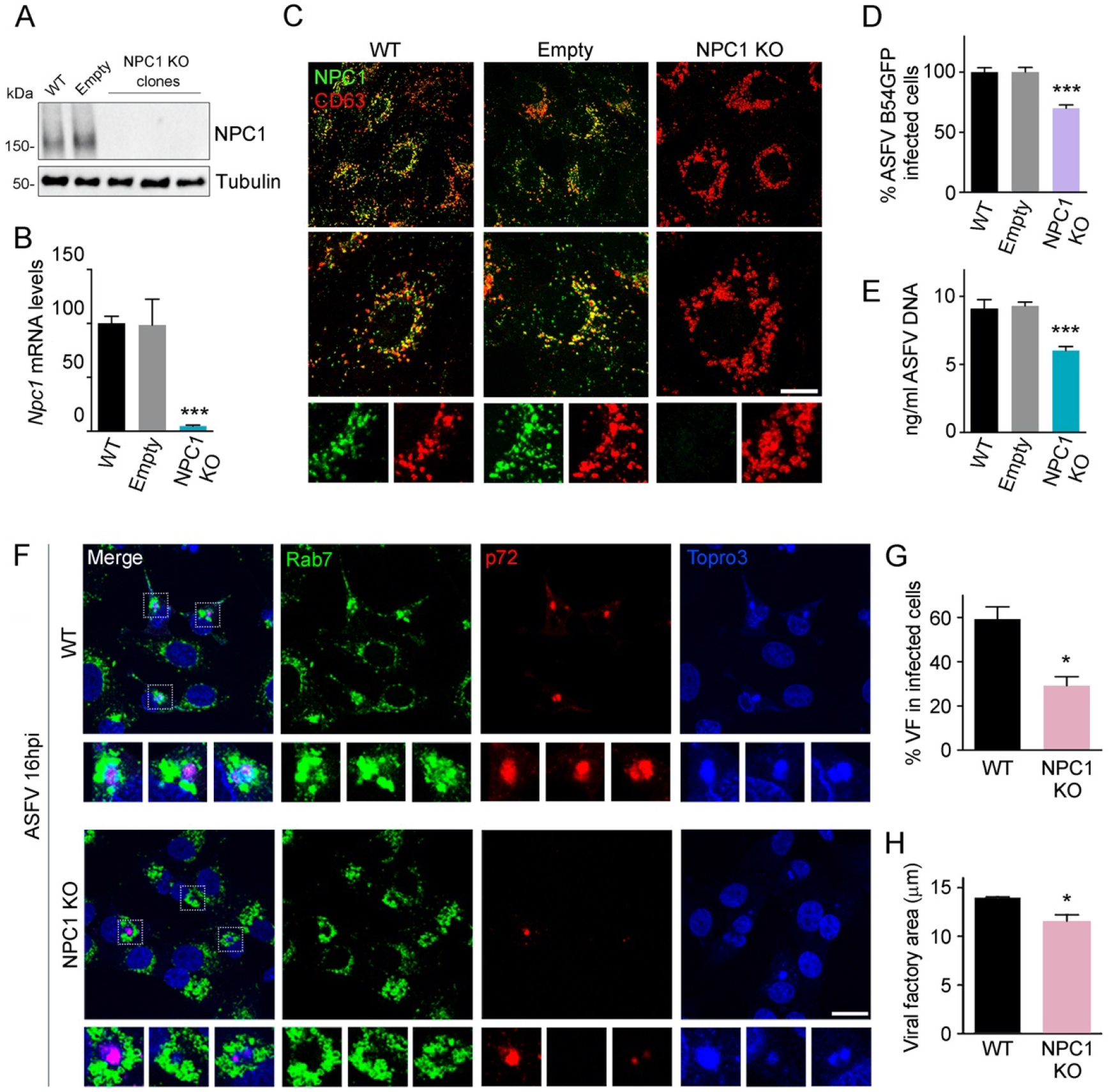
CRISPR KO NPC1 cells phenotype and ASFV infection. (A) Immunoblot of NPC1 and tubulin (loading control) in WT, Empty and several clones of NPC1 KO cells. (B) Npc1 mRNA levels in WT, Empty and NPC1 KO Vero cells detected by qPCR. C) Visualization NPC1 (green) in late endosomes (CD63 (red) in WT, Empty and NPC1 KO cells. Scale bar: 25μm. Zoom images are also shown. (D) B54GFP infection percentages at 16h in WT, Empty and NPC1 KO Vero cells as detected by flow cytometry. Percentages were normalized to values in WT cells. (E) ASFV genome copy number in WT, empty and NPC1 KO ASFV infected cells analysed by real-time PCR. (F) Confocal images of Rab7 (green), ASFV p72 (red) and DNA (Topro3, blue) in WT and NPC1 KO cells. Scale bar: 20μm. Zoom images of ASFV viral factories (boxed regions) are also shown. (G) Percentage of viral factories in WT and NPC1 KO in infected cells shown in F. (H) Quantification of the area of viral factories stained with ASFV p72 in WT and NPC1 KO infected cells shown in F. (B, D, E, G and H). Graphs represent mean±sem from three independent experiments. Statistically significant differences are indicated by asterisks (***p < 0.001, **p < 0.01, *p < 0.05). As controls we used both WT parental Vero cells and cells transduced with empty vector. Once tested that infectivity and replication parameters were the same in both controls, we used Vero WT cells in following experiments.

Then, we analyzed the course of ASFV infection in these validated c14 NPC1 KO Vero cells. A decrease below 40% of infectivity was detected by infection of fluorescent recombinant ASFV, B54GFP by flow cytometry, which is a late protein (Fig 4D). Likewise, ASFV replication was also affected according to decreasing in number of ASFV genome copies (Fig 4E).

Importantly, similar results were obtained by using a different approach by knocking down NPC1 with shRNA (S7 Fig).

The phenotype of NPC1KO Vero cells as expected showed an impediment for cholesterol efflux resulting in the distribution of cholesterol within enlarged endosomes in a closer perinuclear location in aggregates that was absent in control Vero cells (Fig 5A, mock panel).

**Fig 5.**
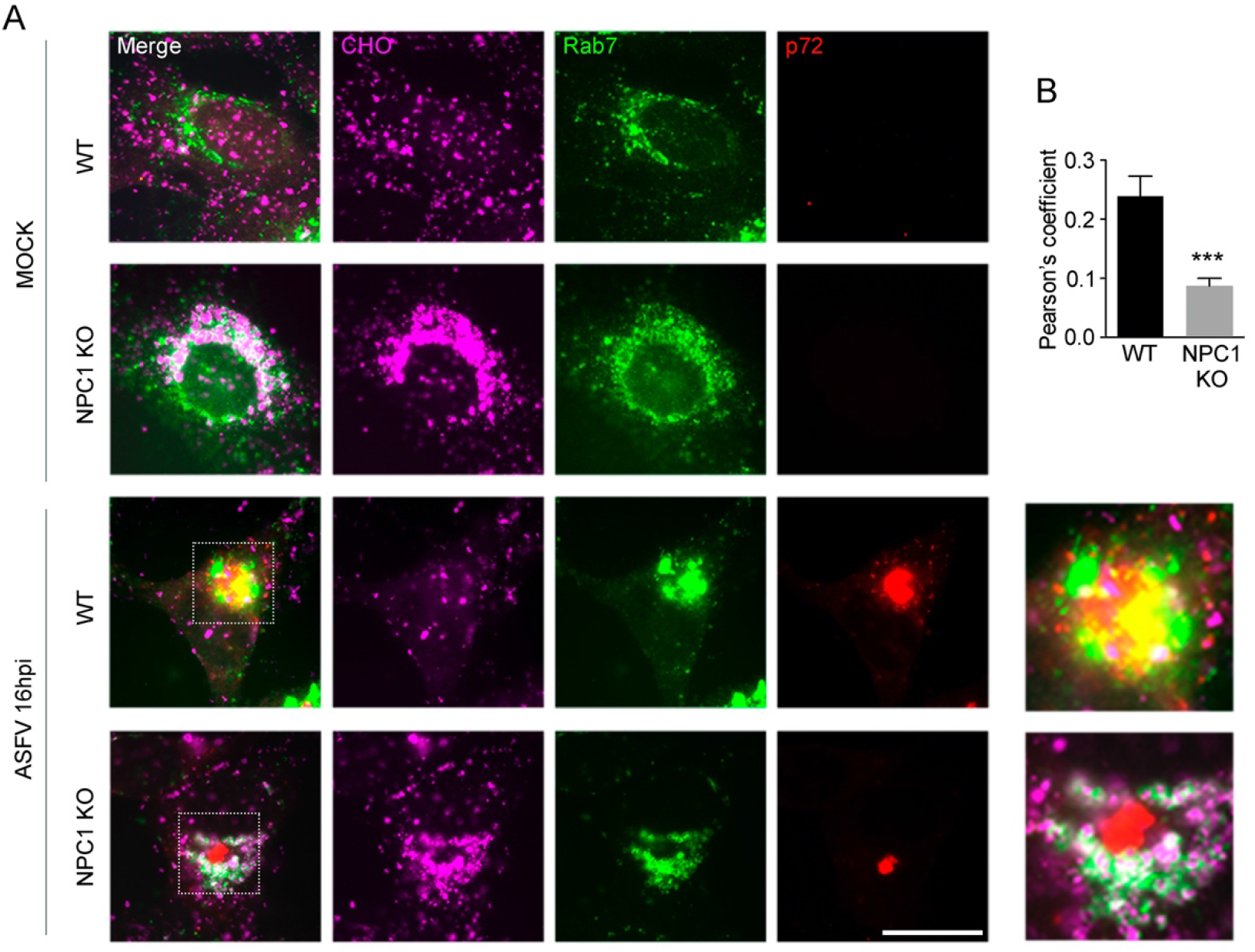
Colocalization of endosomes and viral protein at ASFV factories. (A) Confocal microscopy of merged images (left), cholesterol stained with filipin (CHO, magenta), Rab7 (green) and ASFV p72 (red) in mock or ASFV Ba71V infected Vero WT and NPC1 KO cells. A close apposition and colocalization of endosomes and viral proteins are seen at the factories. Scale bar: 20μm. Zoom images of ASFV viral factories are also shown (right). (B) Quantification of Rab7 and p72 colocalization in WT and NPC1 KO ASFV infected cells using ImageJ software. Graph represents mean±sem of Pearson’s coefficient of three independent experiments. Statistically significant differences are indicated with asterisks (***p < 0.001).

In order to compare the phenotype of mock and infected cells, we used an antibody against ASFV major protein p72 that distributed in VF and accumulated together with viral DNA (Fig 4F, ASFV infected panel). The ASFV factories depicted the characteristic compact appearance in large perinuclear individual aggregates. However, in ASFV infected NPC1 KO Vero cells, the p72 stained VFs were found in a smaller number of cells (Fig 4G) or were reduced in size in a significant number of cells (Fig 4H). In fact, there was less staining with antibody against p72 than DNA staining with Topro3 in infected NPC1 KO Vero cells, probably indicating a reduction in viral protein expression.

Cellular membranes have been postulated as platforms to support and sustain the viral replication site constitution. According to (42), ASFV reorganizes and recruits endosomal membranes forming aggregates and clusters, interspersed and in close contact with the VF. We confirmed this close relationship in infected Vero cells by IFI as a yellow colour was detected in the VFs in the merged image, the viral protein p72 labelled in red, and late endosomal aggregates labelled in green (Fig 5A). Surprisingly, membranes positive for Rab7 in green were also detected around the VF in NPC1 KO cells, but those lacked the usual close contact with the VF as detected by reduction of colocalization percentages, probably suggesting a relevant function for NPC1 in supporting the architecture of the VF (Fig 5B; ASFV 16h panel).

Hence, ASFV infection was altered in NPC1 KO Vero cells, especially important was the unusual architecture of the viral factories and the lower viral infectivity and viral replication with qPCR.

### NPC2 knockdown reduces ASFV infection

To gain mechanistic insight into the ASFV infection cycle, we studied the implication of NPC2 during this process. NPC2 is a single-domain luminal protein that works cooperatively with NPC1 to regulate the egress of endocytosed cholesterol from the late endosome (13, 16, 43–45). To investigate whether NPC2 downregulation affects ASFV infection, we generated lentiviral particles encoding two different shRNA sequences (referred to as 34 and 36) for NPC2 silencing in WT and NPC1 KO Vero cells. The absence of NPC2 expression was confirmed by western blot (Fig 6A). Note that shNPC2-36 sharply decreased NPC2 levels both in WT and NPC1 KO Vero cells, while shNPC2-34 produced only a mild reduction of the target protein. Interestingly, NPC2 levels were significantly increased in NPC1 KO Vero cells as a consequence of NPC1 downregulation, however NPC2 downregulation did not affect NPC1 levels (Fig 6A).

**Fig 6.**
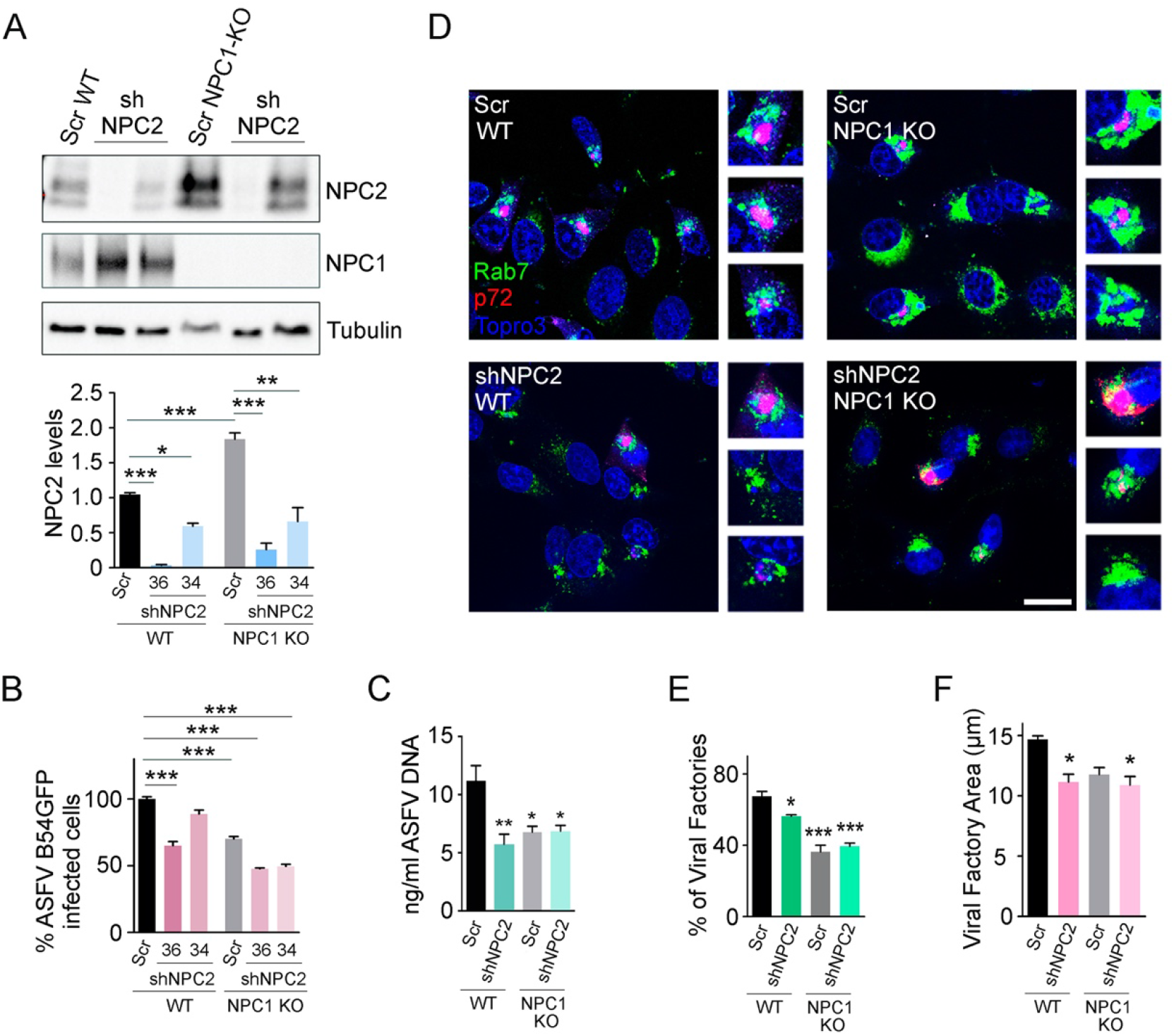
NPC2 downregulation reduced ASFV infectivity. (A) NPC1, NPC2 and tubulin (loading control) were detected by western blot in WT and NPC1-KO Vero cells transduced with Scr (scramble) and shNPC2 (34 and 36) lentiviral particles. Quantification of NPC1 and NPC2 bands were corrected to tubulin data and then normalized to WT-Scr values. (B) B54GFP infection percentages at 16h in WT and NPC1 KO Vero cells transduced with Scr (scrambled) and shNPC2 (34 and 36) detected by flow cytometry. Percentages were normalized to values in Scr-WT cells. (C) ASFV genome copy number in Scr and shNPC2-36 transduced WT and NPC1 KO cells, infected with ASFV and analysed by real-time PCR. (D) Confocal images of Rab7 (green), ASFV p72 (red) and DNA (Topro3, blue) in WT and NPC1 KO cells transduced with Scr and shNPC2-36 lentiviral particles and infected with ASFV. Scale bar: 20μm. Zoom images of ASFV viral factories are also shown. (E) Percentage of viral factories in cells shown in panel E. (F) Quantification of the areas of viral factories stained with ASFV p72 in cells shown in panel D. (A-C, E and F) Graphs represent mean±sem from three independent experiments. Statistically significant differences are indicated y asterisks (***p < 0.001, **p < 0.01, *p < 0.05).

NPC2 downregulation in WT cells showed a reduction of the infectivity measured with B54GFP virus compared to WT-Scr cells. Moreover, in Vero NPC1 KO cells, NPC2 downregulation potentiated the reduction of ASFV infectivity promoted by NPC1 abrogation (Fig 6B). ShNPC2-34 cells showed a discrete phenotype, possibly as a consequence of the mild reduction of NPC2 levels shown in Fig 6A. In this context, we found that ASFV replication was also affected by NPC2 downregulation in WT and NPC1 KO cells, as observed in the quantification of ASFV genome copies by qPCR (Fig 6C).

As we have shown above, NPC1 abrogation altered the distribution and shape of late endosomes and the number and size of VFs in ASFV infected cells. To address whether NPC2 could develop a similar phenotype, we stained Rab7 late endosomes and ASFV p72 protein by immunofluorescence. Downregulation of NPC2 did not induce such an altered phenotype in LE distribution in the factory compared to NPC1 abrogation (Fig 6D); however, the number of ASFV VF and their area were significantly decreased in shNPC2-36 infected cells compared to control cells (Fig 6E and 6F) as occurred in NPC1 KO cells but at moderate levels. Taken together, these results support that ASFV infection was affected by NPC2 downregulation in Vero cells, suggesting an important role of this protein during ASFV infectivity.

## DISCUSSION

ASFV infectious entry relies in the traffic of virions through the endocytic pathway in an orchestrated multi step process depending on several molecular cues that are characteristic of endosomes. Viral uncoating starts with an acidification-dependent stage at the LE, a crucial compartment for the infection (11). Acidic late endosomal environment is able to dissolve the viral capsids composed of major capsid protein p72. Decapsidation allows the subsequent exposure of other internal layers, necessary for further infection events. Beneath the capsid, once the lipid inner viral membrane get exposed, it would eventually merge with the limiting membrane of the LE.

ASFV proteins potentially involved in fusion are located at the inner viral membrane such as candidate fusion proteins E248R and E199L. These have been postulated to be components of a hypothetical complex involved in ASFV fusion, however, no interaction between them could be detected (34). These proteins are highly conserved among ASFV isolates and they share some degree of homology between E248R with VACV fusion protein L1R, and ASFV E199L with VACV A16, A26 and G9 respectively (36). Moreover, E248R and E199L belong to an ortholog cluster of genes from several viruses of the family of nucleocytoplasmic large DNA viruses (NCLDV) (46), sharing conserved structural motifs in several viruses of this broad family, which could indicate a high functional importance of these proteins.

The ASFV fusion process would result in the egress of naked cores towards the cytoplasm in an event that is strongly dependent on cholesterol efflux. Under cholesterol efflux blockade, virions appear retained in endolysosomal vesicles. We have previously shown that impairing cholesterol efflux results in secondary cholesterol accumulation inside endosomes, which is able to inhibit ASFV infection (12). In fact, we found here that both events were impacted at similar concentrations of U18666A compound, which inhibits NPC1 receptor activity. The required concentration of U18666A to inhibit ASFV infectivity was coincident with the necessary concentration to evidence cholesterol retention inside endosomes triggering cholesterol accumulation at the perinuclear area, suggesting that both events are closely related. ASFV infection was blocked upon treatment with U18666A drug, and virions were retained inside lipid-laden endosomes suggesting an alteration at the fusion step.

As a crucial cholesterol transporter protein, this observation also implied a potential role for NPC1 in ASFV infection at the fusion step. Then, we set out to study the viral fusion protein candidates, namely E248R and E199L. These are highly conserved proteins among ASFV isolates located at the inner viral membrane, which get exposed after decapsidation in the LE. Also, E248R has been suggested to be involved in viral membrane fusion and core delivery but not in viral disassembly inside the endosomes. Similarly, E199L is needed for viral core penetration and seems not to be related with later processes, including morphogenesis (33–35). Likewise, proteins belonging to the fusion complex of VACV, a closely related virus, are crucial for DNA replication but not for morphogenesis (37, 47).

In our sequence analysis of ASFV proteins E248R and E199L compared to proteins belonging to the VACV multiprotein fusion complex, we found that E248R yielded 16.2% identity and 30.7% similarity with VACV L1R fusion protein. These interesting sequence similarities would suggest a parallelism between ASFV and VACV fusion proteins and their functions among some of their components.

Also, E248R and L1R have cysteines in similar positions in their sequence, which indicate that the same cysteine oxidation mechanism that generates disulfide bonds on the four cysteines in L1R, could also possibly occur in E248R. These data together with the high sequence conservation of E248R, would suggest a similar function to L1R. In fact, as above explained, we have found here a positive interaction of VACV L1R with NPC1 similar to the ASFV fusion proteins that was previously uncharacterized.

We found a direct interaction between these candidate fusion proteins and cholesterol transporter NPC1 by immunoprecipitation. E248R interacted with NPC1 while no interaction was detected with an independent protein, eIF4e, included as a control in this study. Interestingly, E248R transmembrane domain (TM) was found to be essential in this interaction, since TM domain lacking mutants (ΔTM) were not pulled down by NPC1.

E248R-NPC1 interaction was re-tested and confirmed by reverse pull-down. Besides, individual NPC1 domains A, C and I were included in the analysis, importantly showing that the NPC1 C domain band (along with NPC1) was the only domain detected in the E248R co-transfected bound pellet.

Collectively, these results demonstrate that E248R-NPC1 interaction would occur via E248R TM domain with the NPC1 C domain, as it occurs at EBOV membrane fusion and viral nucleic acid egress from the LE.

ASFV protein E199L was characterized as a virion protein expressed at late times after infection and localized at the virus assembly sites (39). It is a transmembrane protein type I located in the ASFV inner viral membrane. E199L has also been reported to be a positive regulator of the NLRP3-(NLR Family Pyrin Domain Containing 3) and AIM2-(Absent in melanoma 2) inflammasome mediated inflammatory response (48). It has been also reported to be an important antigenic protein in immunoassays (49).

In absence of E199L, ASFV infection drastically decrease and viral particles are retained into LE and lysosomes, being necessary for viral membrane fusion and core penetration (34). Interestingly, E199L resemble weakly in terms of sequence identity to various poxvirus fusion subunits, named VACV proteins A16 (11.1% identity and 20.87% similarity), A26 (10.2% identity and 18.6% similarity) and G9 (12.2% identity and 25% similarity).

Remarkably, likewise E248R, we also found an interaction between E199L and NPC1, confirmed by reverse pull-down. C domain co-transfected with E199L were detected in the immunoprecipitated pellet.

Importantly, NPC1 has been demonstrated as an essential and common partner for successful viral infections including distant viral families. NPC1 is a cholesterol transporter at the endosomal membrane acting as a cellular receptor in the cytoplasmic penetration of Ebola virus (EBOV*; Filoviridae*). Specifically, EBOV protease-primed Glycoprotein (GP) (17) is able to interact with the C domain of NPC1, triggering membrane fusion (50) and the absence of C domain triggers resistance to EBOV infection. The translocation of EBOV GP occurs independently of cholesterol transporter function of NPC1 receptor (19, 51–53). Hepatitis C virus (HCV) (*Flaviviridae*) requires a paralog of NPC1 named Niemann Pick C1 like 1(NPC1L1) (27) on the surface of intestinal enterocytes and human hepatocytes (54, 55), where it is responsible for homeostasis and absorption of cholesterol. Also, a number of reports highlighted the relevance of NPC1 in viral infections like Human immunodeficiency virus type-1 egress (HIV-1; *Retroviridae*). Moreover, the inhibition of NPC1 function by the chemical compound U18666A was reported to inhibit the infection of Chikungunya *(Togaviridae)*, and other flavivirus including Zika Virus (ZIKV), West Nile Virus (WNV), Yellow Fever Virus (YFV) and Dengue Virus (DENV), without any proven interaction yet (26, 30, 32).

Recently, a positive interaction between NPC1 and SARS-CoV-2 Nucleoprotein was reported (22). Consequently, NPC1 appears to be relevant in a variety of RNA or DNA viral families.

Importantly, the U18666A compound is able to block the transport cellular cholesterol from LE to several destinations by binding to the sterol sensing domain (SSD) of NPC1 (19). LE is also the vesicle at which viral membrane fusion occurs. LE is the site of cytoplasmic egress for late penetrating viruses such as ASFV, and its integrity impacts severely viral replication and virion assembly (12). Similarly, entry and replication of DENV, EBOV, ZIKV or CHIKV infections are also profoundly affected under treatment with this drug (32, 40, 56, 57). In fact, compounds specifically designed to interfere with NPC1 C domain, severely impacted ASFV infectivity, supporting our observed interaction between NPC1 C domain and both E248R and E199L (58).

Importantly, NPC1 abrogation completely inhibits EBOV infection, while ASFV as a large complex virus could have evolved alternative cellular pathways, to promote cholesterol efflux homeostasis. From this perspective, it is plausible that due to its genome size, ASFV could have evolved alternative solutions within its more than 150 ORFs, to overcome the abrogation of this cellular gene. However, it is crucial to determine these potential cellular targets to design combined therapies targeting the cellular components required for ASFV infection.

CRISPR KO NPC1 cell lines that inhibit EBOV infection, were partially resistant to ASFV infection while NPC2 knockdown exerted an additive effect. Besides the decrease in infectivity, we observed reduction of ASFV replication in these KO cells. The number of cells harboring viral factories decreased and these presented altered morphology, smaller size and less contacts between the endosomal membranes as supporting elements of the viral factory. Similar to human cytomegalovirus (HCMV; a DNA virus), the constitution of the ASFV viral factory (42), replication and assembly of new progeny occurs in close relationship with lipid-rich cellular membranes. Then, it is possible that an intact cellular cholesterol flux might be also required to facilitate ASFV replication after viral entry.

In summary, ASFV cholesterol efflux blockade in the presence of the U18666A compound drives virion retention inside cholesterol-laden endosomes, inhibiting infection by altering NPC1 function. Our data support a novel NPC1 function to facilitate ASFV membrane fusion and core penetration, according to the direct interaction of ASFV proteins E248R and E199L with this endosomal receptor. Importantly, these ASFV proteins are highly conserved among ASFV isolates and share homology with components of the VACV entry/fusion complex. Binding occurs at the late endosome between the transmembrane domain of both fusion proteins with NPC1 domain C similar to EBOV.

These findings provide further knowledge of the molecular interactions underlying viral fusion and pave the way to the clarification of the function of NPC1 and NPC2 in fusion and early formation of the replication sites for this and other relevant virus models that will be subject of further studies.

## MATERIAL AND METHODS

### Cell lines

Vero and Human embryonic kidney cells (HEK293T or 293T) were grown at 37 °C, 5% CO_2_ atmosphere culture in complete Dulbecco’s Modified Eagle Medium (DMEM) containing 5 or 10% heat-inactivated fetal bovine serum (FBS) respectively, 1% penicillin-streptomycin (P/S) and 1% Glutamax (Gibco, Gaithersburg, MD, USA).

### Viruses and Infection

We used the cell culture-adapted and non-pathogenic ASFV isolate Ba71V (59), the recombinant virus from parental Ba71V, expressing GFP as a fusion protein of p54 protein (B54GFP) (60) isolates or the recombinant Ba71V-30GFP (BPP30GFP) (61). ASFV viral stocks were propagated and titrated by plaque assay in Vero cells, as previously described (59). When using the recombinant virus B54GFP, green fluorescent plaques were observed 4 days after infection under the fluorescence microscope. For immunofluorescence, ASFV stocks were partially purified using a sucrose cushion (40%) in PBS at 68,000 × g for 50 min at 4°C and were further used at a multiplicity of infection (moi) of 1 unless otherwise indicated.

### U18666A chemical compound treatment

U18666A (Sigma) is a chemical compound described as a direct target NPC1 function by binding NPC1 sterol sensing domain (19). It has been widely used to interfere the endosomal cholesterol efflux, mimicking the phenotype observed in Niemann-Pick type C disease (62, 63). Cytotoxicity in Vero cells was previously tested using CellTiter 96 (Promega). Vero cells were pre-treated with DMSO or increasing concentrations of U18666A drug ranging from 0.01-10 μM, 16h before infection without washing leaving compound during the infection.

### Plasmids and constructs

Plasmid pKH3 3xHA was purchased from Addgene (ref. 12555) and used to clone in frame with the HA tag in the N-terminus the viral proteins E248R and E199L into the Bam HI and Eco RI restriction sites.

E248R deletion mutant proteins of distinct domains (Δ constructs) were generated from E248R WT plasmid by site direct mutagenesis using the Q5 mutagenesis kit (New England Biolabs) as follows. E248R ΔExt construct was obtained by deletion of the outer protein domain (from nt 661 to nt 747); E248R ΔTM was obtained by deletion of the transmembrane domain (from nt 601 to nt 660); and E248R ΔExt+TM was obtained by deletion of both TM and Ext domains (from nt 601 to nt 747).

*Sus scrofa* NPC1 WT-linker was generated by gene synthesis (GeneArt, ThermoFisher) with a gly-gly-gly-ser linker followed by 3x Flags at C terminus end as a tag into pCDNA 3.1+.

Plasmids expressing a single domain (A, C or I domain) from full length NPC1 were subcloned as N-terminal 3xFlag fusions into a pCDNA 3.1+ containing the NPC1 signal peptide, the transmembrane domain and the tail. A gly-gly-gly-ser linker was located before the 3xFlag tag.

### Transfections

For pulldowns, 293T cells at 80% confluence were plated 24 hours before transfection in P60 dishes. Prior transfection, the media was changed to DMEM complete media at 2% FCS. Then, the different NPC1 constructs (NPC1, NPC1 A domain, NPC1 C domain and NPC1 I domain) and the viral protein constructs were co-transfected using Lipofectamine 2000 (Invitrogen) following the manufacture instructions.

### Pull downs

293T cells were plated in P60 plates the day before transfection and co-transfected with corresponding pCDNA3.1+ 3XFlag and pKH3 3xHA tagged constructions during 24 hours. Then, cells were lysed in 1ml of ice-chilled lysis buffer consisting in 50 mM Tris, pH 7.4, 150 mM NaCl, 5 mM EDTA, 5% glycerol, 1% Triton X-100 and a protease inhibitor cocktail (complete mini-EDTA free, Sigma), and then, incubated 30 minutes (min) under rotation at 4°C. After, lysates were centrifuged at 15000 rpm for 30 min at 4°C and incubated with Flag coated beads for 3 hours (h) and 30 min at 4°C under rotation. Finally, the lysates were washed five times with wash buffer (50 mM Tris, pH 7.4, 150 mM NaCl, 5 mM EDTA, 5% glycerol, and 0.1% Triton X-100) and bound proteins were eluted in 60 µl of Laemmli buffer and boiled at 100°C. Input and pulled down samples were analyzed by western blot.

### Reverse Pull down for HA viral tagged proteins

Twenty-four hours post-transfection, the cell pellet was resuspended in 200μl of lysis buffer (10mM Tris/Cl pH 7.5; 150mM NaCl; 0.5mM EDTA; 0.5%NP40) and then incubated for 30 minutes on ice. The lysate was then clarified by centrifugation at 14000 x g and diluted five-fold with dilution buffer (10mM Tris/Cl pH 7.5; 150mM NaCl; 0.5mM EDTA) up to 1ml.

The HA pull down was performed using 50μl of the Immobilized Recombinant Protein G Resin (Generon) and specific antibodies against HA (Invitrogen). In order to perform the pull down, a pre-clearing step was done by incubating the diluted cell lysate with 50μl of the Immobilized Recombinant Protein G Resin (Generon) on a rotator for 2 hours at 4°C followed by centrifugation at 2500g for 2 minutes to collect the cell lysate from the beads. Then 2.5μg of the primary antibody was added into the cell lysate and incubated at 4°C on a rotator for another two hours. Then, 50 ul of the protein G resin (Generon) were equilibrated with ice-cold dilution buffer and then incubated at 4°C on a rotator with diluted cell lysate containing the antibody overnight at 4°C on a rotator, followed by centrifugation at 2500g for 2 minutes to remove non-bounds fractions.

The bead pellet was washed once with wash buffer (10mM Tris/Cl pH 7.5; 150mM NaCl; 0.5mM EDTA) and a second time with the wash buffer containing a higher salt concentration (10mM Tris/Cl pH 7.5; 300mM NaCl; 0.5mM EDTA).

After removal of the wash buffer, beads were resuspended in 100μl of Sample Buffer, Laemmli 2X Concentrate (Sigma Aldrich) and boiled at 95°C for ten min to elute the bound proteins. Buffers used for Immunoprecipitations were all supplemented with HaltTM Protease Inhibitor Cocktail EDTA-Free (Thermo Fisher Scientific).

### Western blot analysis and antibodies

Samples from pulldowns were eluted, boiled in Laemmli buffer and resolved by SDS-PAGE in 7%, 12% or 15% acrylamide-bisacrylamide gels or in Mini-PROTEAN TGX Gels (Bio-Rad) then, the gels were transferred to a nitrocellulose membrane (Bio-Rad) using the Trans-Blot Turbo Transfect Pack (Bio-Rad) and the Trans-Blot Turbo system (Bio-Rad) and detected with corresponding antibodies anti-Flag (Sigma), anti-HA (Invitrogen), and anti-Tubulin (Sigma) or HSP90 (Palex) as load controls in Western blot (WB) analysis. As a secondary antibody, anti-mouse IgG (GE Healthcare, Chicago, IL, USA) or anti-rabbit IgG (Bio-Rad) conjugated to horseradish peroxidase was used at a 1:5000 dilution. Finally, bands obtained after development with ECL reagent were detected on a Molecular Imager Chemidoc XRSplus imaging system. Bands were quantified by densitometry and data normalized to control values using Image lab software (Bio-Rad).

### Generation of Vero NPC1-knockout by CRISPR/Cas9 technology

We selected the small guide RNA1 (sgRNA): NPC1 KO Fwd: 5’-CACCGCAAACTTGTATCATTCAGAG-3’ and NPC1 KO Rvs: 5’ AAACCTCTGAATGATACAAGTTTGC-3’, among 4 candidate target sequences at the genomic region of Vero NPC1 region, designed with the tool Deskgen (64). The sgRNA was cloned into LentiCRISPRv2 vector according to the manufacturer indications (GeCKO Lentiviral Crispr toolbox, ZhangLab) and along with plasmids PsPAX2 and VSVg were transfected in 293T cells (as previously described) cultured the day before in p100 plates to generate the lentiviral particles. Similarly, lentiviral particles were also generated with an empty LentiCRISPRv2 as a control (selected clone was named as Empty). Lentiviral particles were harvested 72 h after transfection and centrifugated at 4000 rpm for 3 min to eliminate cellular debris.

The cleared Lentiviral containing medium was transduced in pre-treated Vero cell with Polibrene (8 ug/ml) for 5 min and plated the day before in MW6 plates.

After 72 hours post transduction, the medium was shifted to DMEM 10% FBS containing puromycin (20 ug/ml) to select transduced cells. Finally, Vero cells transduced with lentivirus expressing sgRNA1 were selected after checking the absence of NPC1 by WB and indirect immunofluorescence (IFI) with an antibody against NPC1 (Abcam). We also were able to confirm the typical cholesterol accumulation into laden late endosomes located in a perinuclear position, a usual phenotypic change produced under NPC1 depletion conditions that mimics the Niemann Pick type C disease in CRISPR KO cells. Then, we perform a limited dilution to plate individual clones on 96 MW plate. After clonal expansion during 3 weeks, they were newly seeded in MW6 plates, expanded and finally checked and selected by the absence of NPC1 expression by WB compared to controls, and sequenced with primers Fwd: 5’-ATATATATGAGCGCTCGCGGCCTGG-3’and Rvs: 5’-GCGCGCCTAGAAATTTAGAAGTCGTT-3’. Of the 31 clones examined, we selected c14, c19, c30 and the c14 was used for the experiments (named as NPC1 KO).

### Effect of NPC1 KO on infectivities of pseudotyped VSVs

We further confirmed NPC1 KO cells status by a functional assay testing their resistance to Ebola virus infection using pseudovirions of vesicular stomatitis virus with the EBOV glycoprotein. Infection of control Vero parental cells and Vero NPC1 KO cell lines with recombinant VSV (rVSVs) bearing EBOV glycoprotein (GP) of Mayinga strain or VSV-G were assayed. After 24 h post inoculation, cells were lysed, and luciferase expression measured in a luminometer as Relative Light Units (RLU). The percentages of infectivity were determined by setting the number of RLU in Vero cells to 100% for each envelope.

### Flow cytometry analysis

Detection of ASFV infected cells was performed by flow cytometry. Vero cells were infected with recombinant ASFV B54GFP at a moi of 1 pfu/cell for 16 h. Cells were washed with PBS, harvested by trypsinization, and then washed and collected with flow cytometry buffer (PBS, 0.01% sodium azide, and 0.1% bovine serum albumin). In order to determine the percentage of infected cells per condition, 10,000 cells/time point were scored and analyzed in a FACS Canto II flow cytometer (BDSciences). Infected cell percentages obtained were normalized to values found in control samples.

### Detection and quantitation of the ASFV genome

The quantitation of the number of copies of the ASFV genome was achieved by quantitative real-time PCR (qPCR). DNA from Vero cells infected with ASFV at a moi of 1 pfu/cell for 16 hpi, was purified using the DNAeasy blood and tissue kit (Qiagen) following the manufacturer’s protocol. DNA concentration was measured using a Nanodrop spectrophotometer. The qPCR assay used fluorescent hybridization probes to amplify a region of the p72 viral gene, as described previously (65). The amplification mixture was 200 ng of DNA template added to a final reaction mixture of 20 μl containing 50 pmol sense primers, 50 pmol anti-sense primer, 5 pmol of probe and 10 μl of Premix Ex Taq (2×) (Takara). Positive amplification controls were DNA purified from ASFV virions at different concentrations used as standards. Each sample was included in triplicates and values were normalized to standard positive controls. Reactions were performed using the ABI 7500 Fast Real-Time PCR System (Applied Biosystems) with the following parameters: 94 °C 10 min and 45 cycles of 94 °C for 10 s and 58 °C for 60 s.

For quantitation of *Npc1* mRNA levels, RNA was extracted from Vero cells using RNeasy RNA extraction kit (QIAgen) according to the manufacturer’s protocol. For retrotranscription QuantiTect Reverse Transcription kit (QIAgen) was used to synthesize cDNA, also following the manufacturer’s protocol. 250 ng of cDNA were used as the template for real-time PCR using QUANTITECT SYBR GREEN PCR KIT (QIAgen). Reactions were performed using the ABI 7500 Fast Real-Time PCR System (Applied Biosystems). Expression of *Npc1* gene was normalized to an internal control (18S ribosome subunit), and these values were then normalized to the value of control cells to yield the fold reduction. The following primers were used: *Npc1*_fwd (5′ GTGTGGTGCTACAGAAAACGG 3′), *Npc1*_rev (5′ AAATGCTGCACTGACAGGGT 3′). 18S ribosome subunit primers provided in QuantiTect Primer Assay (Quiagen) were used as internal control.

### Silencing shRNA

Lentiviral vectors containing shRNAs to interfere Npc1 (shNPC1) and Npc2 (shNPC2) were purchased from Merk (Darmstadt, Germany). Two different sequences were transduced for shNPC1 and shNPC2: CCGGGTCCTGGATCGACGATTATTTCTCGAGAAATAATCGTCGATCCAGGA CTTTTTTG (TRCN0000418552, sh NPC1 #52) CCGGAGAGGTACAATTGCGAATATTCTCGAGAATATTCGCAATTGTACCTC TTTTTTTG (TRCN0000421158, sh NPC1 #58) CCGGCGGTTCTGTGGATGGAGTTATCTCGAGATAACTCCATCCACAGAAC CGTTTTTG (TRCN0000293234, shNPC2 #34) and CCGGGCTGAGCAAAGGACAGTCTTACTCGAGTAAGACTGTCCTTTGCTCA GCTTTTTG (TRCN0000293236, shNPC2 #36). A TRC2 pLKO.5-puro Non-Target shRNA was used as control.

Lentiviral suspensions were prepared in HEK293T. HEK293T were transfected with lentiviral and packaging vectors (psPAX2 and p-CMV-VSV-G) using Lipofectamine 2000 ® reagent and OPTI-MEM media (ThermoFisher Scientific, Massachusetts, EEUU) for 4h following the manufacturer instructions. Medium was then changed to Iscove’s Modified Dulbecco’s Medium (IMDM) complemented with 10% FBS, 1% penicillin-streptomycin (P/S) and 1% Glutamax (Gibco, Gaithersburg, MD, USA). Supernatant containing viral particles was collected after 24 and 48h, centrifugated at 3000 rpm for 5 min to eliminate cellular debris and filtered using a 0.45-mm filter.

Vero cells were transduced with lentiviral suspensions directly added to the cells. After 72 hours post transduction, the medium was shifted to DMEM 5% FBS containing puromycin (20 μg/ml) to select transduced cells.

### Indirect immunofluorescence and antibodies, conventional and confocal microscopy

Cells were seeded onto 13 mm glass coverslips in 24 well plates prior to ASFV infection. Then, cells were washed with PBS and fixed with 4% paraformaldehyde (PFA) for 15 min. After a PBS wash, cells were then incubated with 50 mM NH_4_Cl in PBS for 10 min. Then, coverslips were incubated in blocking buffer (0.1% saponin, 0.5% BSA in PBS) for 1 h. Coverslips were then incubated for 1 h in specific primary antibodies diluted in blocking buffer at 37°C. The following rabbit antibodies were used: Rab7 (Cell signaling) and NPC1 (Abcam).

Mouse monoclonal antibodies were: p72 (1BC11, Ingenasa) and CD63 (Novus biologicals). The appropriate secondary antibody conjugated to Alexa Fluor −488 or −594 (ThermoFisher) was used and cell nuclei was detected with TOPRO3 (ThermoFisher). Coverslips were mounted on glass slides using ProLong Gold (ThermoFisher).

Cells were visualized using TCS SPE confocal microscope (Leica) with a 63X Oil immersion objective. The Image acquisition was performed with a Leica Application Suite Advanced Fluorescence Software (LAS AF). All the images were taken with a 1024 × 1024 pixels’ resolution.

To detect free intracellular cholesterol, coverslips were incubated with 100μg/ml filipin (Sigma) for 1h. Filipin signal was recorded using a 390- to 415-nm-wavelength excitation filter and a 450- to 470-nm-wavelength emission filter in a conventional fluorescence microscopy Leica (DM RB). Image analyses were performed with Leica Application Suite advanced fluorescence software (LAS AF) and ImageJ software.

### Sequence comparison

Sequences of pE248R and pE199L proteins were selected from the NCBI data. NCBI reference sequences for proteins of the different ASFV isolates were: txid10498 for Ba71V, for the Georgia strain txid874269 was used, txid561443 for the South African, txid561444 for Namibia and txid561445 for the Kenyan isolates respectively. For Vaccinia Virus (VACV) Western Reserve (WR) proteins A16L, A26L, G9, and L1R, the NCBI reference: txid10254 was used.

BLAST was performed against the E199L protein of ASFV Ba71V, finding 25% identity with A16L fusion protein of Entomopoxvirus Mythimna separata, with NCBI reference: txid1293572. This protein has a 57.3% identity with VACV WR fusion protein A16L.

Alignments between E199L of ASFV Ba71V and A16L of VACV WR, were made using the Geneious 6.0.6 program, with a standard BLOSUM62 matrix as these proteins belong to a different viral family. Sequence similarities were found in the alignments, which could originate a structural similarity and possibly a functional homology to be explored. Alignment with the same matrix was performed between the protein L1R of the VACV fusion complex and the ASFV fusion protein E248R.

Also, alignments of the E248R protein sequences between the different ASFV isolates of were made using the Geneious 6.0.6 program. BLOSUM 90 alignment matrix was used, as this is the most accurate comparison matrix between different isolates of the same viral species. The same was done for the E199L proteins of the different ASFV isolates.

## Statistical analysis

The experimental data were analysed with Unpaired Student *t*-test by Graph Pad Prism 5 software. Values were expressed in graph bars as mean±sem of at least three independent experiments. Statistical significance was assigned at *p<0.05, **p<0.01, ***p<0.001 and ****p<0.0001; n.s, not significant.

## Acknowledgements

This research was partially supported through “La Caixa” Banking Foundation award number LCF/PR/HR19/52160012, Spain; Spanish Ministry of Science and Innovation, Spain RTI2018-097305-R-I00) Instituto de Salud Carlos III, ISCIII, Spain FIS PI 1801007 and the European Commission, Horizon 2020 Framework Programme European Union ASFVInt-ERANET-2021-0017 and VIRUSCAN FETPROACT-2016-731868.

## Supporting information

**S1 Fig. E248R and E199L structure resembles to VACV fusion machinery**. Comparison of the amino acid protein sequences of E248R (ASFV, Ba71V isolate) with L1R (VACV WR); and E199L (ASFV Ba71V isolate) with their potential equivalents in VACV A16, G9 and A26. The identical amino acids are indicated in green or purple respectively, in grey the amino acids conserved between 60 and 80%, and the amino acids conserved less than 60% in white.

**S2 Fig: E248R and mutants, E199L binding to NPC1.** Membranes used to compose figure 3B. Dashed boxes were taken to create the western blot composition showed in the figure. Membranes were revealed with mouse anti-Flag antibody, mouse anti-HA antibody and mouse anti-tubulin.

**S3 Fig. VACV L1R binding to NPC1.** (A) Membranes used to compose figure 3C and (B) 3D. Dashed boxes were taken to create the western blot composition showed in the figure. Membranes were revealed with mouse anti-Flag antibody, mouse anti-HA antibody and mouse anti-tubulin.

**S4 Fig. Reverse Pull down E248R.** Membranes used to compose figure 3F. Dashed boxes were taken to create the western blot composition showed in the figure. Membranes were revealed with mouse anti-Flag antibody, mouse anti-HA antibody and mouse anti-tubulin.

**S5 Fig. Reverse Pull down E199L.** Membranes used to compose figure 3G. Dashed boxes were taken to create the western blot composition showed in the figure. Membranes were revealed with mouse anti-Flag antibody, mouse anti-HA antibody and rat anti HSP90.

**S6 Fig. NPC1 KO validation in Vero cells.** (A) NPC1 detection in Vero or Vero NPC1 KO cell lines (B) Indirect immunofluorescence depicts NPC1 in green detected with a specific antibody against NPC1. (C-D) NPC1 KO cells depicts dilated endosomes detected in red with the acidic probe Lysotracker. Cholesterol stained with Filipin III (in blue) accumulates in dilated vesicles, similarly as occurs in cells pre-treated with U18666A drug. (E) Infection of Vero and NPC1-KO-Vero cells with recombinant VSV (rVSVs) pseudotyped with Ebolavirus Glycoprotein (EBOV-GP) Mayinga strain or VSV-G 24 h. Cells were lysed 24 hs post-infection and assayed for luciferase expression. Percentages of infectivity were determined by setting the number of RLU in Vero cells to 100% for each envelope.

**S7 Fig. NPC1 KD impact ASFV infectivity.** (A) NPC1 detection in Vero WT Scr (scrambled) cells or Vero NPC1 KD (shNPC1-52) cells with a rabbit anti-NPC1 antibody. (B) B54GFP infection percentages at 16h in Vero WT Scr and Vero NPC1 KD cells (shNPC1-52) as detected by flow cytometry. (C) ASFV genome copy number in Scr and Vero NPC1 KD (shNPC1-52), infected with ASFV and analysed by real-time PCR.

## Notes

### Competing Interest Statement

The authors have declared no competing interest.

